# Fixational eye movements depend on task and target

**DOI:** 10.1101/2021.04.14.439841

**Authors:** Norick R. Bowers, Josselin Gautier, Samantha Lin, Austin Roorda

## Abstract

Human fixational eye movements are so small and precise that they require high-speed, accurate tools to fully reveal their properties and functional roles. Where the fixated image lands on the retina and how it moves for different levels of visually demanding tasks is the subject of the current study. An Adaptive Optics Scanning Laser Ophthalmoscope (AOSLO) was used to image, track and present Maltese cross, disk, concentric circles, Vernier and tumbling-E letter fixation targets to healthy subjects. During these different passive (static) or active (discriminating) fixation tasks under natural eye motion, the landing position of the target on the retina was tracked in space and time over the retinal image directly. We computed both the eye motion and the exact trajectory of the fixated target’s motion over the retina. We confirmed that compared to passive fixation, active tasks elicited a partial inhibition of microsaccades, leading to longer drifts periods compensated by larger corrective saccades. Consequently the fixation stability during active tasks was larger overall than during passive tasks. The preferred retinal locus of fixation was the same for each task and did not coincide with the location of the peak cone density.

## Introduction

When fixating our gaze on an object, our eyes are never truly at rest. Even while staring at a small object, like the bottom row of a Snellen acuity chart, our eyes are constantly in motion. Small, fast microsaccades and slow drifts constantly shift the image of the fixation target over the photoreceptor lattice. It is becoming increasingly clear that these small movements are not simply noise in the oculomotor system. Recent studies have shown that fixational eye motion serve a number of functions in the visual system, such as preventing fading (Martinez-Conde, Macknik, Troncoso, & Dyar, 2006), reallocating attention within the fovea (Hafed & Clark, 2002; Engbert & Kliegl, 2003; Ko, Poletti, & Rucci, 2010; Poletti, Listorti, & Rucci, 2013; Intoy & Rucci, 2020), enhancing the discrimination of fine spatial details through a combination of spatiotemporal enhancement at high frequencies (i.e. whitening of the power spectrum) (Rucci, Iovin, Poletti, & Santini, 2007; Rucci, 2008) and enrichment of the information relayed from the retina to the brain through dynamic sampling (Anderson, Ratnam, Roorda, & Olshausen, 2020; Ratnam, Domdei, Harmening, & Roorda, 2017; Burak, Rokni, Meister, & Sompolinsky, 2010).

A limitation in studying the smallest fixational eye movements (FEM) is the instrument used to measure the gaze itself (Poletti & Rucci, 2016). Modern video-based eye trackers are convenient but often lack the resolution of earlier systems. Scleral search coils, which were among the first high-resolution eye trackers, have high temporal and spatial resolution (Robinson, 1963; Collewijn, Van Der Mark, & Jansen, 1975) but are invasive and difficult to use. Dual Purkinje image (DPI) eye trackers came to use shortly after search coils and offered a noninvasive way to attain high accuracy for tracking the gaze (Crane & Steele, 1985) and still represent a reliable eye tracker today (Fourward Technologies, Gallatin, MO). However the DPI system has been shown to have its own drawbacks: Oscillation and prismatic effects of the crystalline lens can give rise to spurious measurements of gaze direction (Tabernero & Artal, 2014; Nyström, Andersson, Magnusson, Pansell, & Hooge, 2015; Bowers, Boehm, & Roorda, 2019). On the other hand, modern video eyetrackers, today’s most commonly used instruments in research and industry, suffer from pupil size changes. (Choe, Blake, & Lee, 2016; Nyström, Hooge, & Andersson, 2016; Hooge, Holmqvist, & Nyström, 2016). They also simply lack the requisite spatial resolution to accurately estimate the gaze position produced by these fixational eye movements (Kimmel, Mammo, & Newsome, 2012; Holmqvist & Blignaut, 2020).

FEM have seen a resurgence of interest in research lately as their role in fine-scale vision has been reexamined. The effects of fixation target and task have not been thoroughly examined using high resolution retinal-image-based tracking techniques. Published studies, which primarily relied on video eyetrackers, showed generally that smaller targets elicit modestly less overall FEM compared with larger targets (McCamy, Jazi, Otero-Millan, Macknik, & Martinez-Conde, 2013; Kazunori, Kana, Risako, Wakana, & Nobuyuki, 2016). Other target properties, such as shape, color, contrast, blur and luminance in eliciting improved fixation have been more scarce (Thaler, Schütz, Goodale, & Gegenfurtner, 2013; Steinman, 1965; Bhattarai, Suheimat, Lambert, & Atchison, 2019; Ukwade & Bedell, 1993) with few trends emerging except that bull’s eye and cross targets (or a combination of both) elicit the least FEM (Thaler et al., 2013). Most experiments that require fixation use a simple static fixation target (see Thaler, 2013 for a comprehensive overview of the variety of targets commonly used).

Although some studies have shown that FEM can have implications in encoding visual information and can be modulated during fine-scale fixation tasks (Rucci & Victor, 2015; Martinez-Conde et al., 2006), little is known about how active fixation tasks might be used to elicit an overall more stable fixation. A more comprehensive characterization of FEM and fixation target is therefore important for two reasons. First, it will continue to build our knowledge on the remarkable ability for fine foveal vision in humans. A second, more practical, reason is to learn what target and/or fixation task might minimize overall FEM in clinical settings where motion and its consequent blurred or distorted retinal images can be detrimental, such as with fundus photography, OCT scans or microperimetry.

The current study aims to compare and contrast FEM during *active* fixation tasks - those that contain temporal variation and require subject input - and *passive* fixation tasks, where the subject is simply instructed to maintain fixation on a target. An Adaptive Optics Scanning Laser Ophthalmoscope (AOSLO) is used as an eye tracker to acquire high spatial (<1 arcmin) and temporal (960 Hz) resolution eye traces. Since the AOSLO can also obtain an unambiguous record of the motion of the target that is projected onto the retinal surface, we compare how the preferred retinal locus for fixation (PRL) relates to the location of peak cone density (PCD) for each type of fixation target.

## Methods

Eight healthy subjects, 3 male and 5 female, with normal or corrected-to-normal vision participated in the experiment. Subject ages ranged from 23-53 years old. All experimental procedures adhered to the conditions set by the institutional review board of the University of California, Berkeley and followed the tenets of the Declaration of Helsinki. Each subject read and signed a written informed consent document. Prior to imaging, the subjects’ eyes were dilated and cyclopleged using 1 drop each of 1% tropicamide and 2.5% phenylephrine. The drops were used to provide maximum dilation for imaging as well as to paralyze accommodation, both of which help to ensure high quality images in the AOSLO. No detectable difference has been found between eye traces measured in an SLO system with or without dilation (Bowers et al., 2019).

### AOSLO System

Data were recorded using the Adaptive Optics Scanning Laser Ophthalmoscope (AOSLO) (Roorda et al., 2002), which is used to image and track the retina as well as to provide the fixation targets used in this experiment. For imaging, a point source of light is relayed through the optical path and scanned across the retina in a raster pattern utilizing two scanners, a 16kHz fast horizontal scan and a 30Hz slow vertical scan. The reflected light is descanned through the optical path and directed through a confocal pinhole to a custom-built Shack-Hartmann wavefront sensor and a photomultiplier tube (Hamamatsu, Japan). The Shack-Hartmann wavefront sensor is used to measure the optical aberrations and send a correction to the deformable mirror (7.2mm diameter, 97 actuators membrane; ALPAO, Montbonnot-Saint-Martin, France) in the optical path. Light detected by the PMT and the positional information from the scanner are combined to construct videos of the retina with 512×512 pixel sampling resolution at a frame rate of 30Hz (the speed of the slow vertical scanner). For this experiment, the imaging wavelength was 680nm, with 940nm used for wavefront sensing. The field size of the video was 0.9 × 0.9*◦*. Using an average power of 50-70 *μW*, the raster scan field appeared as a bright red square to the subject. Fixation targets were presented to the subject within the red field by turning off the scanning laser using an acousto-optic modulator (Brimrose Corp, MD) at the appropriate time points during the raster scan. To the subject, these targets appeared as black-on-red decrements. The stimuli were very sharp and had high contrast owing to the use of adaptive optics on the input scanning beam. Importantly, these decrements are also encoded directly into the video, which allows for an unambiguous measurement of the motion of the image of the fixation target over the retina. This system is capable of obtaining near diffraction-limited images of the photoreceptor mosaic and delivering stimuli with the precision of 1 pixel (∼6 arcseconds). An example video from one of the trials in this experiment is shown in the Supplementary Materials. This system has been explained in greater detail in previous manuscripts from our group. (Poonja, Patel, Henry, & Roorda, 2005; Rossi & Roorda, 2010).

### Experiment Design

The experiment consisted of 5 different conditions: Maltese cross, disk, concentric circles, Vernier acuity, and a tumbling E (M, D, C, V, and E respectively). The Maltese cross condition (M) was chosen as it has been suggested to provide a better fixation target than the simple dot that is commonly used in fixation tasks. The disk condition (D) consisted of an annulus within the center of the raster that the subjects were instructed to fixate. Both of these conditions were simple passive fixation tasks where subjects were instructed to hold their gaze on the target. The concentric circles condition (C) consisted of concentric rings moving in a constricting radial motion. There were 6 rings ranging in size from 10 to 1 arcmin that were presented over the course 18 frames (3 frames per ring size) and replayed every 30 frames for a frequency of 1 Hz. The aim of the concentric rings was to provide a simple fixation task that was more visually engaging than a static target. The Vernier hyperacuity condition (V) required subjects to judge the relative displacement of two tiny horizontal bars which appeared at random intervals (seven 6-arcsecond steps). The tumbling E condition (E) consisted of a tumbling E task where the subjects were asked to report the orientation of a letter E as it rotated randomly. The size of the E varied in seven steps, from 20/6 to 20/20 Snellen acuity. For both V and E tasks the stimulus was presented for 0.5 sec (15 consecutive frames) and there were random time intervals between presentations - evenly spread over 0.5 to 1.5 sec - where nothing was presented. The random time intervals were used so that subjects could not anticipate the next trial and were therefore compelled to maintain fixation the entire time. The V and E condition can be differentiated from the others as they both required active subject judgment and response, as well as providing temporal variation. These conditions were further categorized into passive tasks (M, D) and active tasks (E, V) for further analysis depending on whether they required subject response and varied in time. The concentric circles provided a mixed task as it had temporal variations but did not require subject response. See Table 1 for the differences between the conditions. The different fixation targets were presented in a pseudo-random order to eliminate any training or fatigue effects. Furthermore, subjects were given consistent instructions from a script to avoid known changes in behavior due to instruction (Steinman, Cunitz, Timberlake, & Herman, 1967). The full script is provided in the Supplementary Materials but the primary emphasis in the instruction was for the subject to maintain their gaze throughout the entire duration of each 30-second trial task. There were five 30-second trials for each condition in total.

**Table 1:**
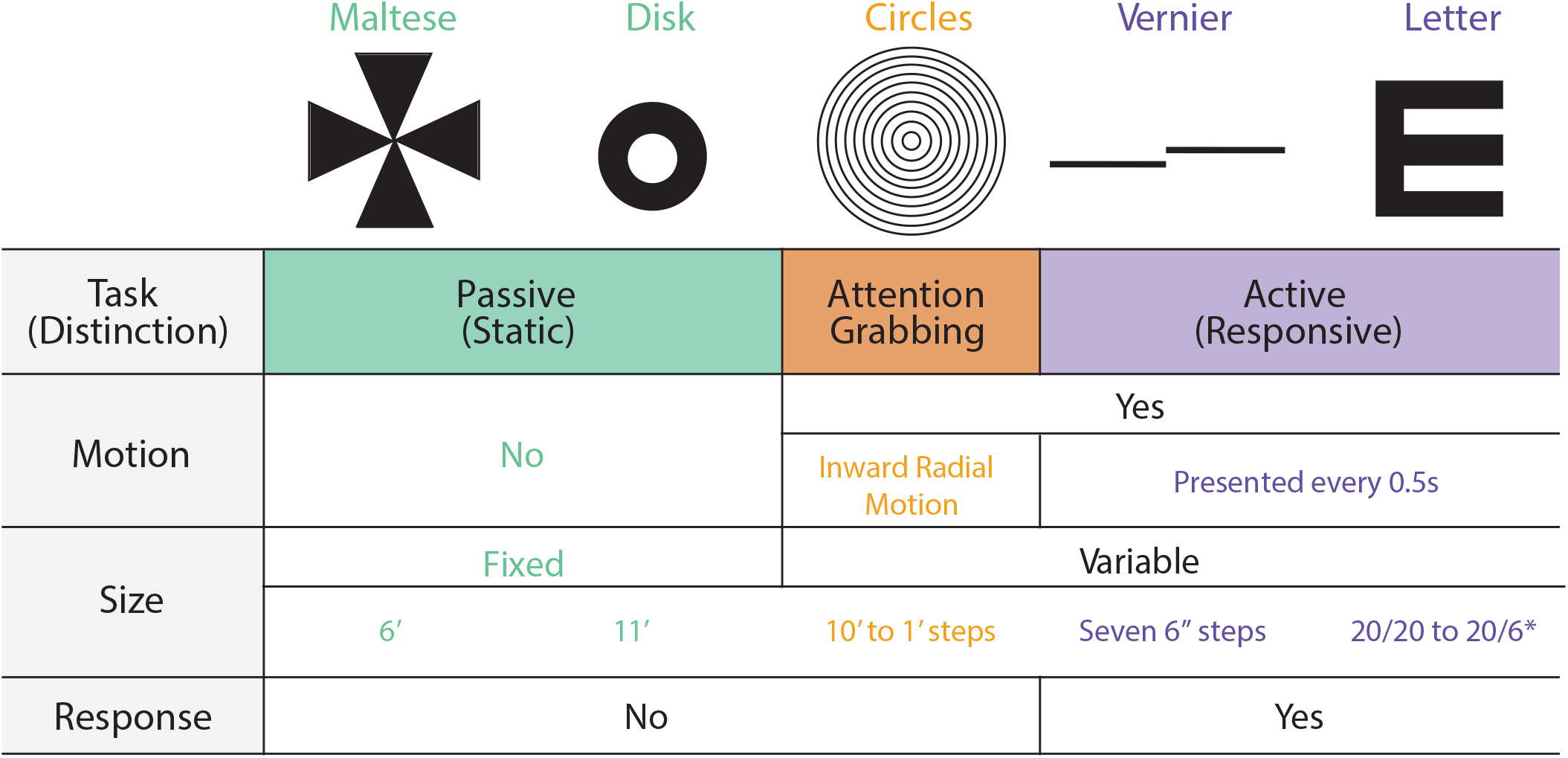
Illustration of the 5 experimental conditions and their respective parameters. The different colors indicate distinctions between passive (green), active (purple), and mixed (orange) tasks. This color scheme will be used throughout to differentiate the 5 conditions. *Snellen size

### Eye Tracking and Video Processing

Since this system utilizes a raster scanning technique (i.e. each frame is acquired over time), additional eye motion information is available beyond the 30Hz frame rate. This additional information can be extracted to achieve eye traces at temporal resolution many times greater than the frame rate of the movies (Stevenson & Roorda, 2005; Vogel, Arathorn, Roorda, & Parker, 2006). In order to acquire eye traces at higher temporal resolution than the 30Hz framerate, each frame of the AOSLO movie is broken into horizontal strips and cross-correlated against a reference frame. This analysis is done offline using custom software written in Matlab (The Mathworks, Natick, MA) (Stevenson, Roorda, & Kumar, 2010). This technique allows collection of eye traces at high spatial (<1 arcmin) and temporal (960Hz) resolution. Eye traces were separated into drifts and saccades using a semi-automatic software and the output was manually verified by the authors. Saccade onset was defined as the point when instantaneous speed exceeded 1.5deg/sec and offset was defined as the point when the trace fell back below this threshold. Blinks were defined as frames of the AOSLO movie when the mean luminance fell below a threshold that was defined on a per-subject basis dependent on the average brightness of the respective movies.

The AOSLO records high-resolution videos of the retina for each trial and the fixation target is directly encoded into the video, thereby making it possible to plot the exact path of the fixation target over the photoreceptor mosaic directly. This is done in the following way. First, the eye motion traces extracted from AOSLO videos indicate how the entire retina moves, but do not directly indicate where the fixation target lands on the retina. Computing the actual retinal trajectory requires computing the ∆X and ∆Y offsets that need to be applied to each eye trace to anchor it to the exact position of the fixation target on the retina. To accomplish this step, we first generate a high quality master retinal image chosen from one of the best videos recorded in the experimental session for each subject. Then we use the same cross correlation methods to align strips containing the encoded stimulus from each video with that master retinal image and determine the position of the stimulus on the master retinal image. The X-Y position corresponding to the strip that contains the stimulus is then aligned to that exact position on the master retinal image using these ∆X and ∆Y offsets. In theory, the offset only needs to be computed once for a single strip, but the match between a single strip and the master retinal image can have small errors due to noise in the strip or torsion in the retinal image. So, to improve accuracy, we compute the average offsets from at least 20 unique strips, ensuring that the standard deviation of the offsets is less than 2 pixels (0.2 arcmin). These processing steps yield accurate trajectories in retinal coordinates for every trial and every condition, all referenced to a single master retinal image.

For all of our subjects the master retinal image was of sufficient quality to label all cones across the image. Cones were labeled across the entire foveal region using a combination of automatic cone-finding (Li & Roorda, 2007) with manual intervention when necessary. Cone density was computed within a 10-arcmin diameter circular window while it traversed, pixel by pixel, across the mosaic (using a convolution process). The 10-arcmin averaging window was chosen since it has been shown to strike an optimal balance between minimizing noise and maximizing resolution (Wang et al., 2019). The point of maximum cone density was expressed as the pixel location with the highest density value. This analysis allows us to determine how the location on the retina the subject used to examine the stimulus (the Preferred Retinal Locus of fixation, or PRL) differed from the peak cone density (PCD) on the retinal lattice. Figure 1 shows an example of a master retinal image from one subject with selected structural and functional measures overlaid onto it.

**Figure 1:**
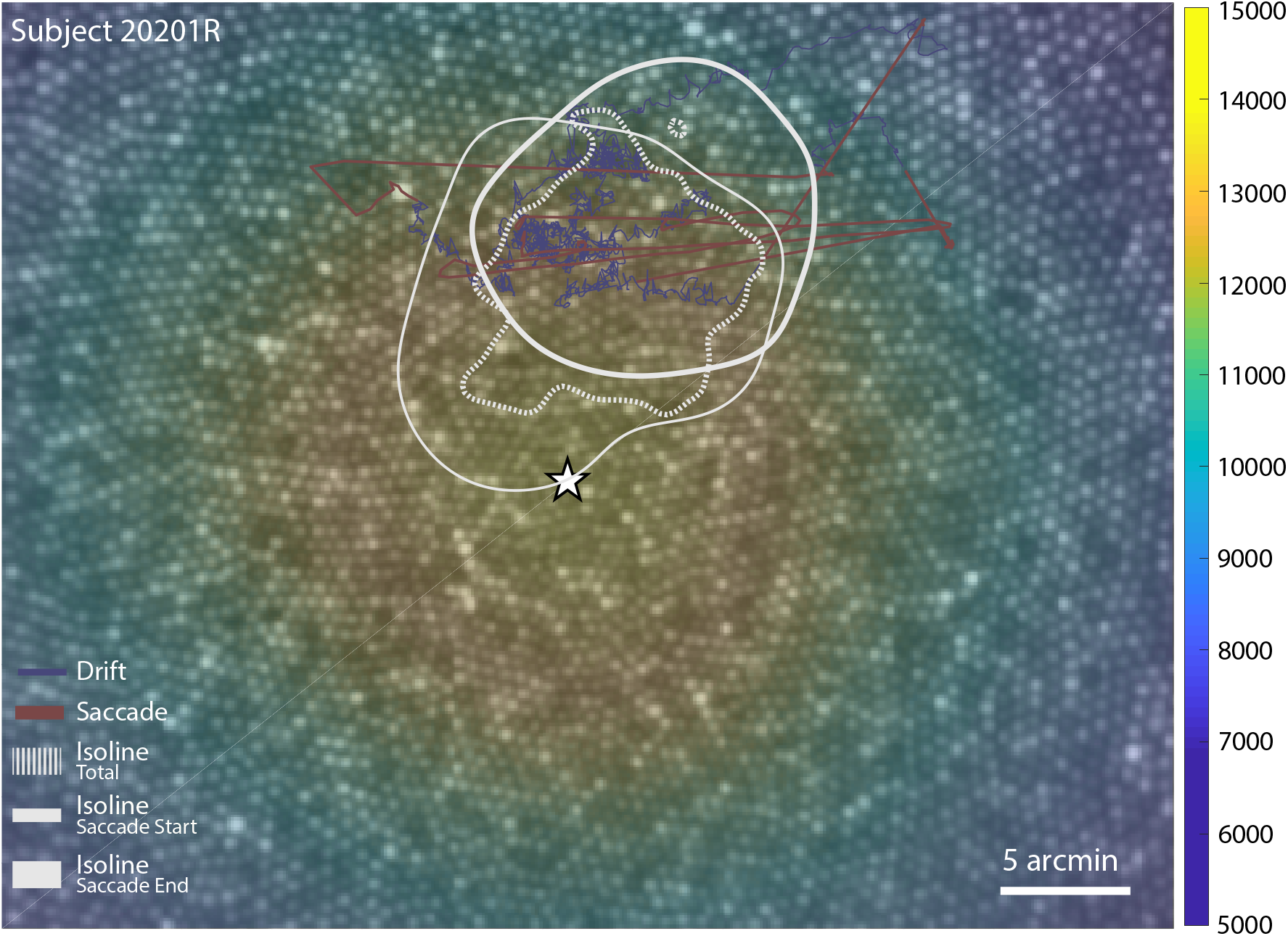
Master retinal image for subject 20201R with functional and structural measures overlaid. The star indicates the point of maximum cone density (PCD) and the underlying colormap represents the cone density in cones per degree^2^. 5 seconds of eye movement is plotted on the retinal image representing the stimulus motion on the retina with saccades (red) and drifts (blue) highlighted. The isoline contours for all the eye positions obtained during the Vernier condition (dotted), as well as saccade start (thin) and end points (thick) for this condition are shown in gray (see Figure 5 and Figure 7 for contours for all conditions and all subjects).

### Eye Movements, ISOA and PRL Analysis

The Isoline Area (ISOA) method to measure fixation stability was proposed as a better alternative to Bivariate Contour Ellipse Area in the presence of multiple loci of fixation positions (Castet & Crossland, 2012). It does not make any assumption on the nature of the random variables underlying the distribution of data points, which is specifically appropriate for people with eccentric fixation (Whittaker, Budd, & Cummings, 1988), but also normal subjects whose fixational eye positions have been shown to be not randomly distributed (Cherici, Kuang, Poletti, & Rucci, 2012). The ISOA and PRL are computed through Kernel Density Estimation (KDE) of the 2D probability density function (PDF) of eye positions. The ISOA is the area within the non-uniform contour that encompasses 68% of the entire eye trace. The PRL is computed as the corresponding peak of the 2D PDF. In other words, the isoline contour encloses all the eye positions that lie within 1 SD from the PRL, if we could assume normality and a unique PRL (which we observed). In order to assert both non-normality of the 2D distribution of eye positions and the non-separability between pairwise distributions obtained during different conditions, one-way and two-way 2D Kolmogorov-Smirnov tests were used, respectively. This implementation relies on Fasano & Franceschini’s generalization (Fasano & Franceschini, 1987) for two dimensions.

## Results

### Global eye movement statistics

FEM under the different conditions are plotted in Figures 2-4. Figure 2 reveals expected extensive differences in FEM between subjects (Cherici et al., 2012). The intersubject variability will be discussed later, but in order to better examine differences between conditions, the data were normalized (All the non-normalized data for each subject and each condition is in the Supplementary Materials). Figure 2 illustrates how this was accomplished using microsaccade amplitude as an example. First, every FEM measurement was collected for each subject in each condition (top, left). Then each subject’s own measurements were normalized so that each subject’s mean performance across all 5 conditions (bottom, left) was equal to unity. Then, all subjects’ data were averaged together to compute normalized performance across conditions (right). Tasks were separated into three categories depending on the extent of active participation required of the subject. Passive tasks required the subject to simply maintain fixation on a stationary target. The passive tasks are the Maltese cross and the disk, since both of these required the subject to hold their gaze on an unchanging target. Active tasks were those in which the subject had to respond to a changing target. The active tasks consist of the Vernier and the tumbling E conditions. The concentric circles condition represents a mixed task because, while it had temporal variations as the circles moved inward, there was no response required of the subject.

**Figure 2:**
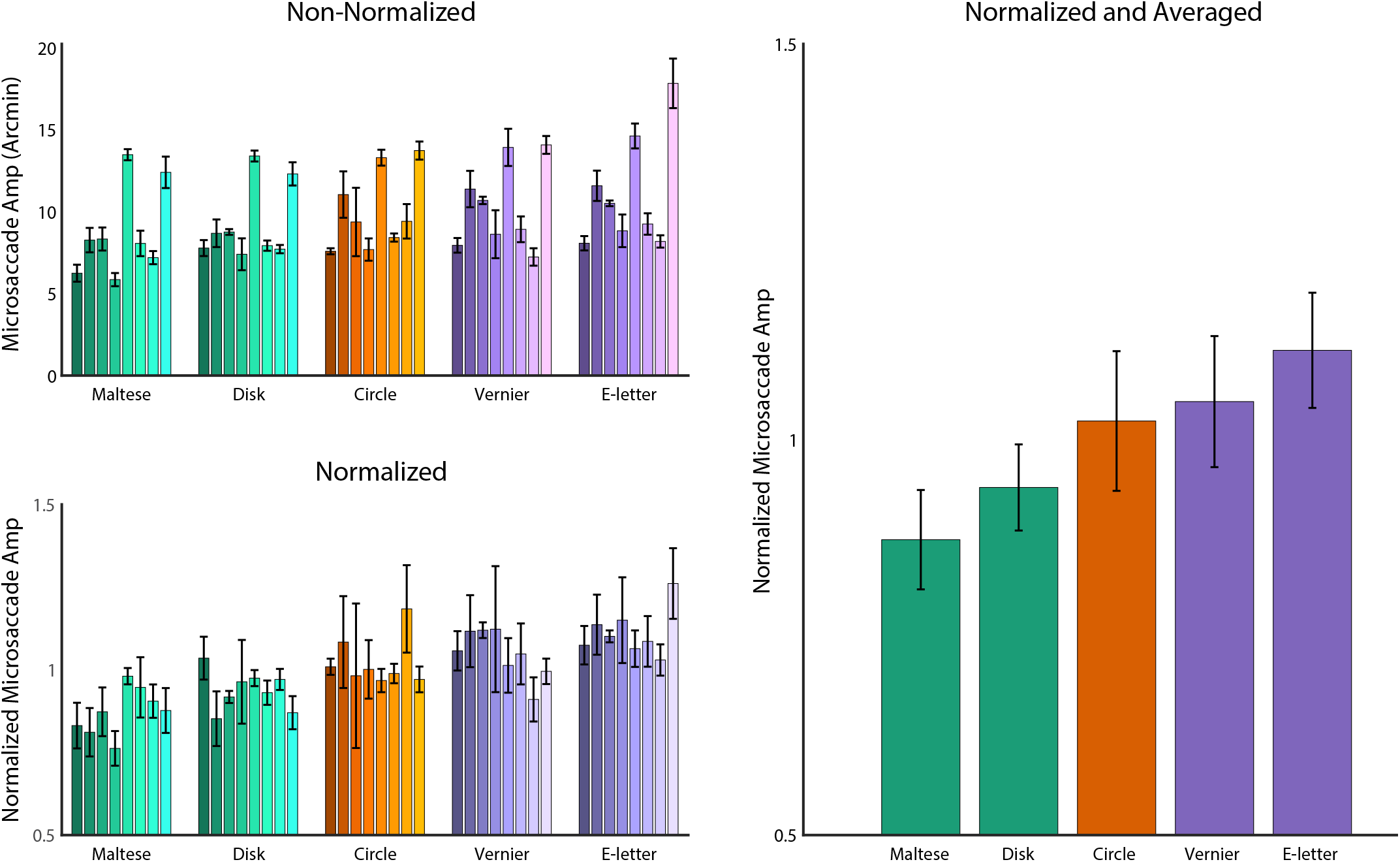
Illustration of the normalization process for each metric using microsaccade amplitude as the example. The different colors represent further distinction between passive, active, and mixed tasks used in later analysis. **Top, Left:** Every subject’s individual measurement for this parameter. Different shading indicates different subjects. **Bottom, Left:** Normalized measurements across conditions. The measurement for each subject was normalized to the mean across all 5 conditions. **Right:** Normalized performance across all conditions. The colors represent the passive (green), active (purple) and mixed (orange) tasks. Error bars represent S.E.M.

Selected FEM measurements are plotted in Figure 3. Overall, during the active tasks the subjects drifted for longer periods of time and made fewer microsaccades. The amplitude of saccades were larger in the active tasks and the corresponding saccade rate was significantly lower. The drift amplitudes and duration were also correspondingly larger in the active tasks. This is in line with previous research showing that subjects will suppress their microsaccades while performing high-acuity visual tasks (Rucci et al., 2007). Subjects would suppress their microsaccades, drift away from the target, and then be forced to make relatively larger microsaccades to reorient themselves during the active discrimination tasks. The active tasks also had overall larger fixation as indicated by the isoline area. Overall, these data reveal that there was more motion in the active tasks vs the passive tasks, even though there were fewer saccades.

**Figure 3:**
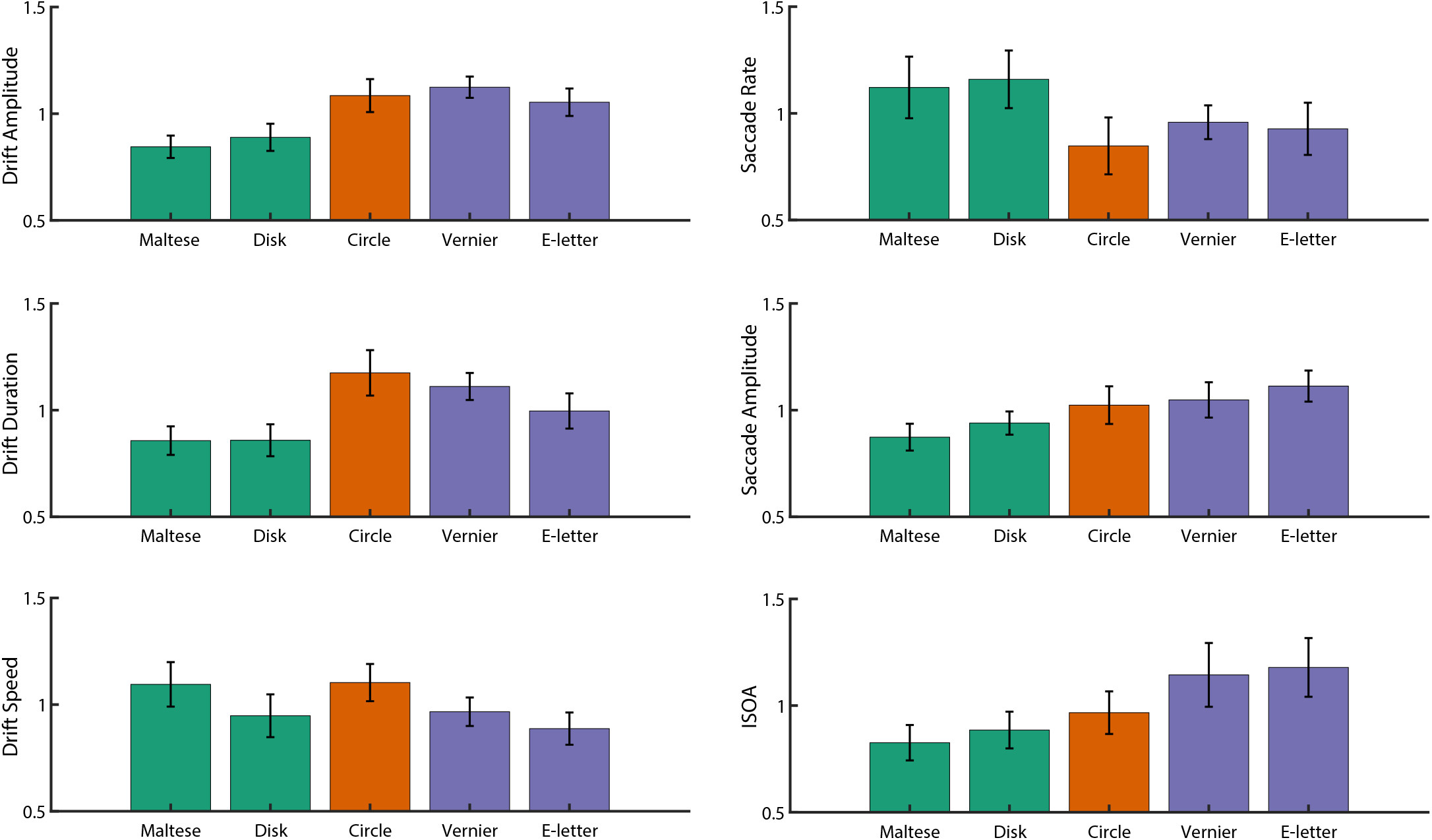
FEM measurements across the five conditions. In general, the eye tended to have longer and larger intersaccadic periods of drift in the active fixation tasks where subject response was required (purple bars). Whereas during the passive fixation tasks (green bars), subjects tended to make shorter but more frequent saccades. The circle task (orange bar) represents a mixed task. While it did not require subject response, it still had temporal variations that were significant enough to differentiate it from a simple fixation task. Subject behavior in the circles tasks did not readily align with either the passive or the active tasks. Error bars represent S.E.M.

In recent years, fixation has been considered a more “active” process than previously thought (Rucci & Victor, 2015). In order to tease apart the effects of different fixation targets, it is prudent to analyze behavior under active fixation conditions, in which the subject is required to attend and respond to a changing target; compared to passive fixation tasks, where the subject is required to simply hold their gaze on a stationary target. The circle was excluded from this analysis since it represents a mix of these two distinct categories, and behavior during the circle task did not readily align with either the passive or active tasks. Since the two active tasks (Vernier and Tumbling E) and the two passive tasks (Maltese and Disk) showed similar patterns of behavior relative to each other (One-way ANOVA (F(4,35)<0.001, post-hoc Tukey-Kramer, p>0.05 for all cases of active and passive), they were further combined to examine the differences in active fixation vs passive fixation, regardless of the task involved. The combined tasks are plotted in Figure 4. The trend of fewer, but larger microsaccades is more readily evident when comparing active and passive tasks pooled together.

**Figure 4:**
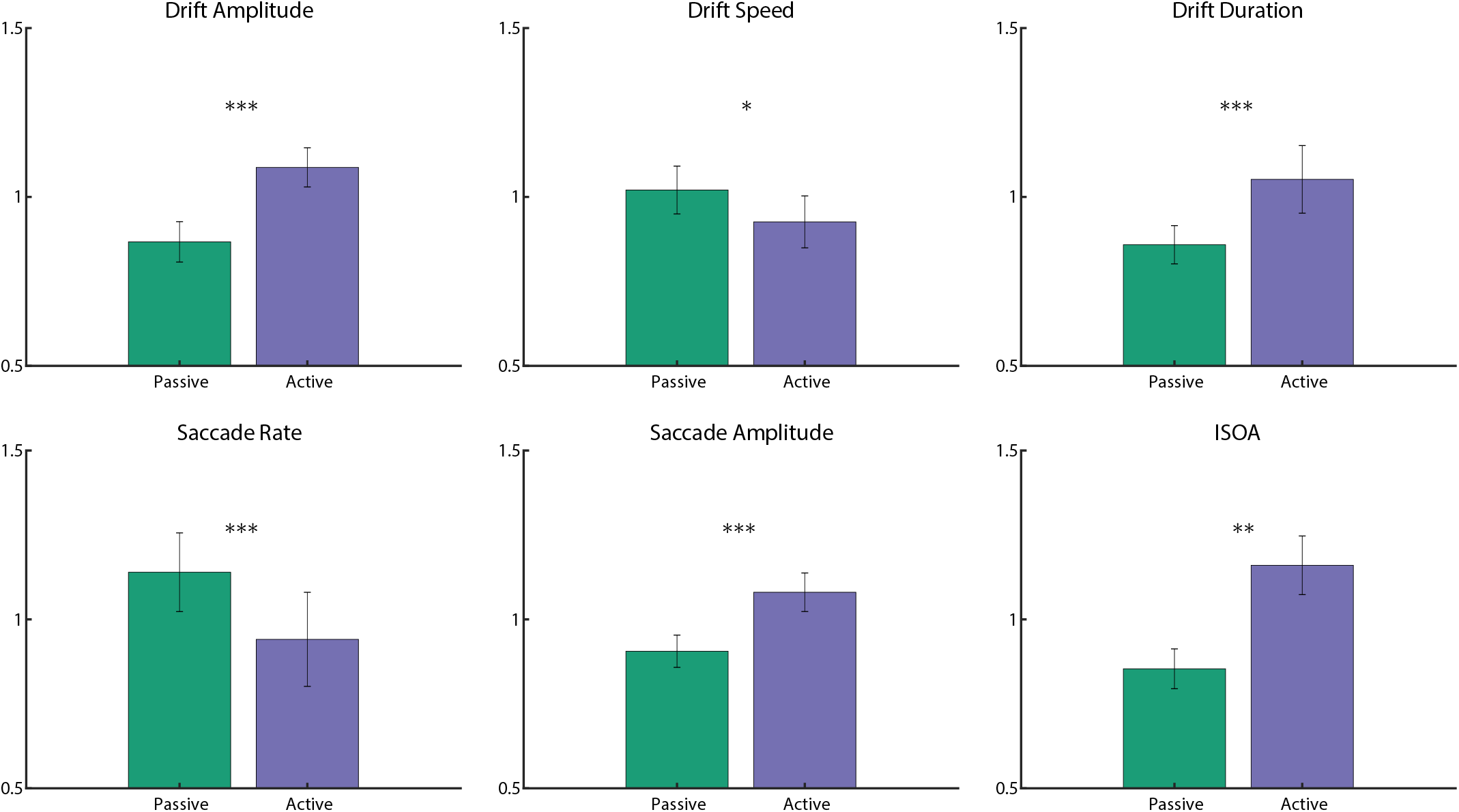
After the data was collapsed into active and passive tasks (excluding the circle task), the same variables were tested together. Active and passive tasks were averaged together, as well as their error bars in order to simplify the analysis to just the active vs the passive tasks. Error bars represent S.E.M and the asterisks represent levels of significance (repeated-measures t-test, p<0.05, p<0.01, and p<0.001 respectively).

Figure 5 shows the 68% isoline contours for each subject for each condition centered on the PRL, which is defined at the peak of the fixation positions’ PDF. The stimulus for each condition is drawn in the center of each graph for reference, but the stimulus will sweep across the retina based on the extent of the eye motion. Although the overall fixation area was larger for the active tasks compared to the passive tasks (two-sampled t-test, p<0.01). Extensive intersubject variability is readily apparent in Figure 5. The average standard deviation of the ISOA between subjects for each condition (columns in Figure 5) was 43.53 arcmin^2^. Whereas the average standard deviation between conditions for each subject (rows in Figure 5) was 21.26 arcmin^2^. Although intersubject variability is extensive (roughly double the size of the difference between conditions), there is still a significant difference in fixation behavior between the five conditions. Large differences in fixation behavior between subjects is expected in measurements of fixational eye movement, especially when psychophysical expertise is taken into account (Cherici et al., 2012).

**Figure 5:**
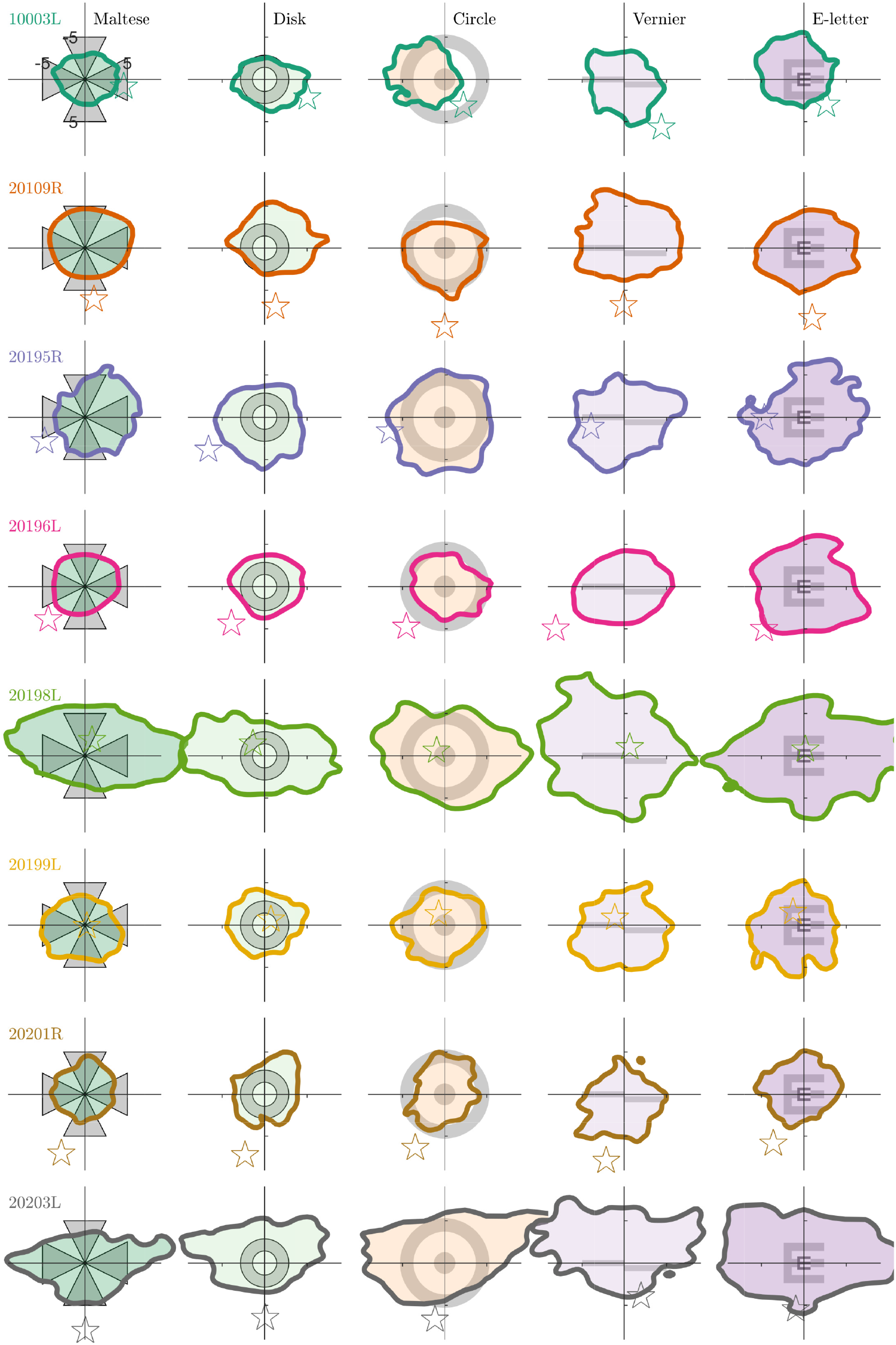
68% isoline contours (ISOA) of the entire eye movement trace. Each row is a subject and each column is a fixation condition, the stimulus for which is drawn centered on each plot. Position (0,0) on the plot corresponds to the PRL, or the peak of the fixation position distribution. Note that idiosyncrasies in the eye movement can cause this peak to appear displaced from the center of the isoline contour. The position of these distributions represent the location of the fixated image and are plotted in fundus-view coordinates (same as Figure 1). The star in each plot indicates the relative location of the PCD. Axis units in the upper left plot are in minutes of arc.

Figure 6 shows the same data as shown in Figure 5 but in this case the ISOAs from all subjects and conditions are overlaid on a single plot with all subjects’ respective PCD at (0,0). This figure reveals several phenomena. First, the PRL rarely coincides with the PCD and is displaced, on average by 5.20 arcmin (SD=2.54 across tasks and between subjects, SD = 0.23 across subjects and between tasks). This is largely in line with other reports that the PRL does not perfectly correspond with the PCD (Putnam et al., 2005; Li, Tiruveedhula, & Roorda, 2010; Wilk et al., 2017; Wang et al., 2019). Second, the PRL tends to be displaced above the PCD in fundus coordinates. This is consistent with recently published reports (Reiniger, Domdei, Linden, Holz, & Harmening, 2019). Finally, subjects adopt a consistent PRL regardless of the task and its visual demand. The Euclidean distance between the PRL and the peak cone density did not significantly differ from one another (repeated measures ANOVA with a Greenhouse-Geisser correction (F(2.155, 15.087) = 0.313, P = 0.751). A Kolmogorov-Smirnov test for two-dimensional distributions was used to determine whether eye position distributions, and therefore the PRL, differed between the different conditions. There was no difference in the PRL location between conditions for any subject (p-value <0.001 in each of the 5 conditions and 8 subjects). This is in agreement and extends on the finding that the PRL for a static Maltese cross target remains stable between days (Kilpeläinen, Putnam, Ratnam, & Roorda, 2020).

**Figure 6:**
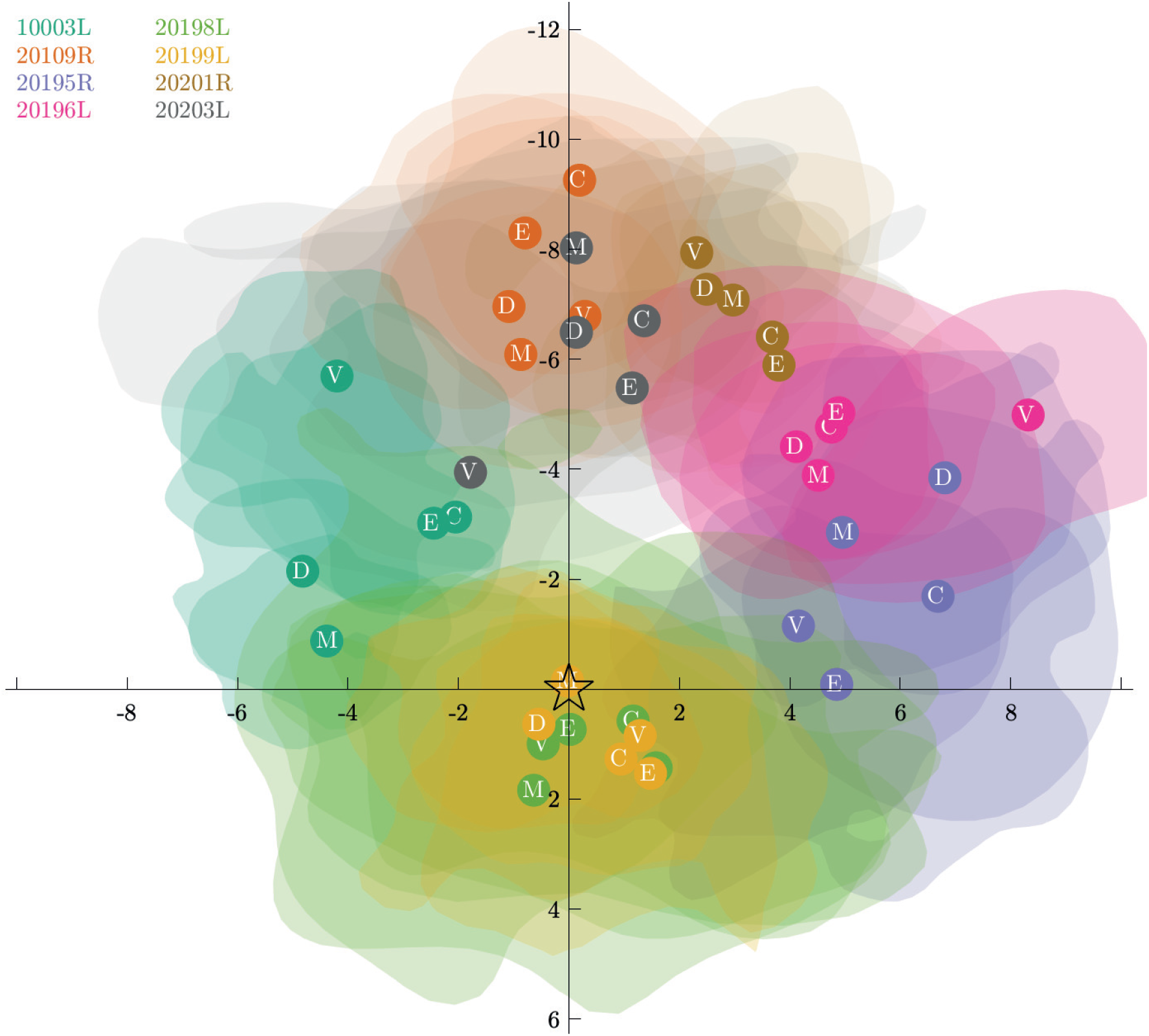
Individuals’ PRL locations plotted relative to their PCD centered to (0,0). Location of the PRL’s are shown as opaque circles overlaid onto the transparent isoline contours with the condition defined in white text within the center. This figure highlights that each subject’s PRL tends to fall off their respective PCDs. However, the various PRLs defined in each condition all group together closely. To enhance visibility, the isoline contours here encompass only 38% (0.5 S.D.) of the fixation trace instead of the 68% (1 S.D.) used in the rest of the study. Axis units are in minutes of arc.

Figure 7 shows 68% isoline contours for saccade start and end points, represented by thin and thick contours respectively. Some clear and distinct patterns emerge here. First, the ISOA for saccade start and end points cover a larger area than the conventional ISOA. Second, the distribution of saccade landing positions appears smaller than at their initiation positions, especially for subjects 20109R, 20195R and 20196L. Third, the distribution of microsaccades tend to be horizontally spread, which is largely in agreement with other research (Cherici et al., 2012; Sheehy et al., 2020; Thaler et al., 2013). Finally, despite the a more extensive horizontal spread, every subject shows a tendency to make saccades, on average, in an upward direction. The saccades move the image upward in the fundus, straddling either side of the PRL. If the image moves up during a saccade, this means that the fovea moves down relative to it. In gaze coordinates this corresponds to a saccade that redirects the gaze upward as the coordinates between fundus view and gaze coordinates are inverted. This movement could be classified as a form of spontaneous upbeat micro-nystagmus (Eggers et al., 2019) although in these instances, the upbeat nystagmus clearly does not indicate a pathological condition. A similar behavior is reported in other papers (Mestre, Gautier, Bedell, Douton, & Pujol, 2019; Stevenson, Sheehy, & Roorda, 2016), but not observed universally, perhaps owing to differences in how the saccades were plotted.

**Figure 7:**
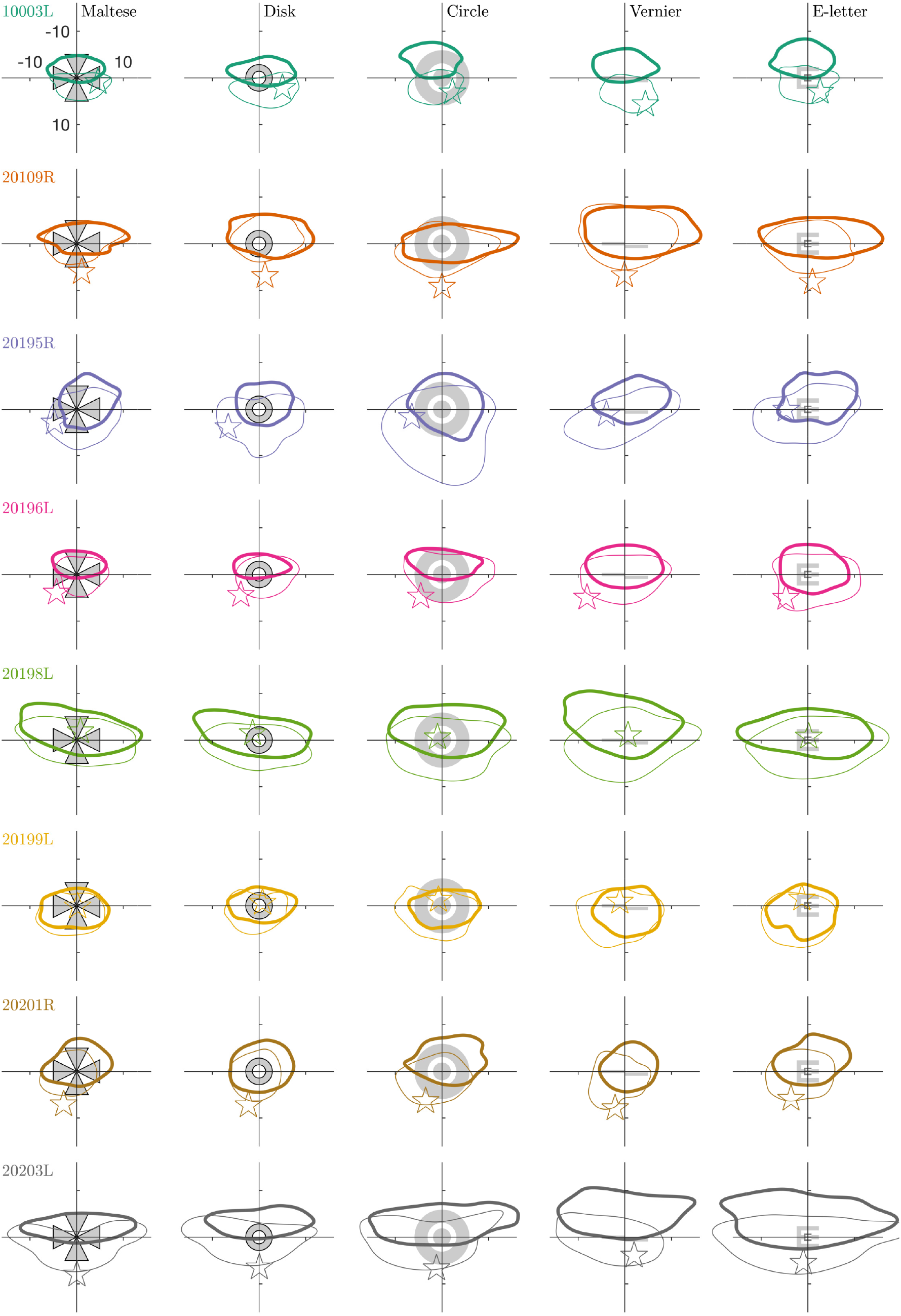
Isoline contours for the location of the fixated image at the start and end of saccades, drawn as thin and thick contours respectively. Each row is a subject and each column is a fixation condition, the stimulus for which is also drawn on the figure. Position (0,0) on the plot corresponds to the PRL location from Figure 5. The star indicates the location of the PCD. Axis units in the upper left plot are in minutes of arc.

## Discussion

Our study has shown that large and significant variations of microsaccade and drift kinematics exists between subjects and between different tasks. We have confirmed that an individual’s FEM behavior depends on the task involved. Specifically, we have found that active tasks result in less frequent microsaccades and correspondingly longer and larger drift epochs which, in turn, cause the microsaccades to be larger. This pattern of behavior leads to overall larger ISOAs during active fixation tasks. Passive tasks, by comparison, are marked by shorter and more frequent microsaccades and smaller, briefer drift epochs, all leading to a smaller ISOA. The increase in ISOA from passive to active tasks was 57% (+/− 23% S.D.) on average. These results can be explained in part by previous findings that suggest that FEM is an active behavior that is subconsciously mediated to serve different functions depending on the task at hand (Rucci & Victor, 2015). During the active tasks subjects tended to suppress their microsaccades because the rapid transients from these movements can be detrimental to fine-scale discrimination due to microsaccadic suppression, either from blurring of the retinal image or central suppression.

The stability of the PRL was tested using a 2-dimensional Kolmonorov-Smirnov test and was found to remain the same regardless of the fixation target. This extends on a recent report that the PRL for a Maltese cross target does not change between hours and across days (Kilpeläinen et al., 2020). However, the possible shifts in the location of the PRL have not been investigated for binocular viewing conditions or for more complex viewing experiences, such as during smooth pursuit or fixation within extended scenes.

Although the peak cone density on the retina offers the best location for photoreceptor spatial sampling (according to the Nyquist sampling limit), the PRL very rarely aligns with it. Previous research has consistently shown the same (Putnam et al., 2005; Wang et al., 2019; Wilk et al., 2017). We found the average separation between the PCD and the PRL to be 5.20 arcmin. In the majority of cases the PRL is positioned superior to the PCD (in fundus-view coordinates). This reflects a similar tendency for the PRL in individuals with a central scotoma to adopt an eccentric PRL in the superior retina (Messias et al., 2007; Verdina et al., 2017). This means that the retinal location with peak cone density is sampling a part of the visual field just above the direction of gaze. This tendency has been reported previously by another group (Reiniger et al., 2019) who also used an AOSLO. These displacements are very small and in our opinion, as discussed in one of our previous papers (Wang et al., 2019), do not have any functional importance.

A subclinical form of upbeat nystagmus was present to varying extents in all of our subjects. Similar behavior was reported for some, but not all of the subjects in two other studies (Stevenson et al., 2016; Mestre et al., 2019) and is further evident in a slight upward tendency (although not commented on) in the saccade distribution plots of other papers (Cherici et al., 2012; Thaler et al., 2013). Interestingly, we found that these corrective saccades appear to only partially “refoveate” the stimuli (i.e. reposition the image on the PRL), instead they generally landed above it. This suggests that the visual system might be very much aware of its own oculomotor ability and could be anticipating the likely direction of the following drift so that the target will fall on the PRL during the middle of the drift segment. In any case, the minutiae of FEM reveals that a PRL that is identified by any of the current methods, including the ISOA approach used here, may be ill-defined. This is a topic of an ongoing investigation.

While we measured significant and informative differences in FEM between conditions, we found that differences in FEM between individuals are even greater. The standard deviation of the ISOA between individuals, for example, was roughly twice that between conditions. These differences can be partly explained by experience (Cherici et al., 2012). All of our subjects were recruited from within the UC Berkeley School of Optometry community and therefore had some experience sitting for visual psychophysics experiments and/or for clinical examinations. But subjects 10003, 20109 and 20196, who all had ISOAs that were lower than the mean, have logged dozens of hours in psychophysics experiments related to eye tracking, including AOSLO psychophysics experiments.

When considering these small fixational eye movements it is prudent to consider the goals for maintaining a subject’s steady fixation, since all fixations are not equal. If the goal of the fixation target is to minimize the overall movement of the eyes (as is the case in many clinical situations), then one must consider which types of FEM are most likely to be an impediment. If the rapid transients from saccades are most likely to have a deleterious effect then it is preferred to rely on an active fixation task so the subject will suppress their microsaccades in order to perform the task. If the goal is to minimize the total area covered by the fixation, then choosing a more passive fixation target is likely to be most effective. Finally, if the goal is to measure FEM and understand their role in regards to ecologically valid situations, it is recommended to use active fixation tasks, since these tasks will reflect real-world discrimination of fine stimuli.

## Conclusion

This study examined the influence of different fixation targets and tasks on FEM and the location of the PRL in healthy eyes. Using an AOSLO, we developed a new method to locate and follow the target projected on the retina over time relative to the PCD. We confirmed the non-normality of the eye motion distribution, hence the necessity to rely on better descriptors of fixation stability indices such as ISOA and its accuracy to estimate each individual’s PRL. The different fixation tasks consisted of active tasks, which had temporal variation and required subject responses, and passive tasks, where the subjects were instructed to simply hold their gaze on the target. The active tasks elicited larger but fewer microsaccades. Consequently, the amplitude and duration of intersaccadic drifts were significantly larger. Larger and longer drifts combined with larger microsaccades led to larger overall fixation instability, as quantified by the ISOA. Our result suggests that subjects surpress their microsaccades during active tasks, and the subsequent longer drift epochs would cause the object to move away from the PRL, thereby requiring a relatively larger microsaccade to reorient. Finally, although the FEM were significantly modulated by the task, the intersubject variability was expectantly substantial. The two-to-four times larger effect on fixation stability across individuals compared to task suggest that experimenters might, when aiming to better control the user’s eye position, put a greater emphasis on instructions, training, and subject recruitment rather than on the fixation stimulus itself.

## Acknowledgments

Authors would like to acknowledge William Tuten, Swati Bhargava, Pavan Tiruveedhula and Ally Boehm for their support in collecting data. Supported by National Institutes of Health grants: R01EY023591, T32EY007043 and T35EY007139.

## Disclosures

AR has a patent (USPTO7118216) assigned to the University of Houston and the University of Rochester which is currently licensed to Boston Micromachines Corp (Watertown, MA, USA). Both he and the company stand to gain financially from the publication of these results. No other authors have competing interests.

© 2009–2010, Tobias Elze.

## Supplementary Materials

### Example Video

The attached movie is a 5 second segment from one trial of the concentric rings condition for subject 20201R. The movement of the retina arising from fixational eye motion and its amplitude relative to the size of the stimulus is readily evident. This segment also highlights how the decrement stimulus is inscribed into the video file directly, providing a completely unambiguous record of the stimulus’ position across the photoreceptor mosaic. Note that this segment has been compressed to keep the file size small and does not represent the raw videos used in stabilization and analysis.

### Instructions

The instructions given to each subject were identical from one subject to the next. The below script was read out to each subject for each condition and any clarifying questions were answered before the experiment began. Note that experimenters asked the subject to blink when their tear film degraded to the point of interfering with the video quality.

Maltese: “In this experiment you will be required to look towards the center of the cross for the entire duration of the 36 second video. There will be a sound played when the video starts and stops recording. You are allowed to blink as needed and I will ask you to blink if necessary. Press the start button when you are ready to go.”

Disc: “In this experiment you will be required to look towards the center of the disc for the entire duration of the 36 second video. There will be a sound played when the video starts and stops recording. You are allowed to blink as needed and I will ask you to blink if necessary. Press the start button when you are ready to go.” Concentric circles: “In this experiment you will be required to look towards the center of the target for the entire duration of the 36 second video. You will see a series of rings that become smaller. There will be a sound played when the video starts and stops recording. You are allowed to blink as needed and I will ask you to blink if necessary. Press the start button when you are ready to go.”

Vernier: “In this experiment you will be required to look at the two lines for the entire duration of the 36 second video. The two lines will vary in position at random intervals which will be announced by an audible cue. Using the up and down buttons, report if the right line is higher or lower than the left line. There will be a sound played when the video starts and stops recording. You are allowed to blink as needed and I will ask you to blink if necessary. Press the start button when you are ready to go.”

Tumbling E: “In this experiment you will be required to look at the “E” for the entire duration of the 36 second video. The letter “E” will change size and direction at random intervals which will be announced by an audible cue. Using the buttons, report the direction the “E” is facing: up, down, left, or right. There will be a sound played when the video starts and stops recording. You are allowed to blink as needed and I will ask you to blink if necessary. Press the start button when you are ready to go.”

### Extra tables

The attached .csv files contain the raw, unnormalized data for each of the 6 eye motion parameters shown in Figures 3 and 4. This is the same data shown in the top, left of Figure 2, but for each parameter. Rows indicate different subjects and columns indicate different conditions. The variable names and respective units are shown in the top, left of each table.

## Notes

### Competing Interest Statement

The authors have declared no competing interest.

